# Site-specific amino acid distributions follow a universal shape

**DOI:** 10.1101/2020.08.05.238493

**Authors:** Mackenzie M. Johnson, Claus O. Wilke

## Abstract

In many applications of evolutionary inference, a model of protein evolution needs to be fitted to the amino acid variation at individual sites in a multiple sequence alignment. Most existing models fall into one of two extremes: Either they provide a coarse-grained description that lacks biophysical realism (e.g. *dN/dS* models), or they require a large number of parameters to be fitted (e.g. mutation–selection models). Here, we ask whether a middle ground is possible: Can we obtain a realistic description of site-specific amino acid frequencies while severely restricting the number of free parameters in the model? We show that a distribution with a single free parameter can accurately capture the variation in amino acid frequency at most sites in an alignment, as long as we are willing to restrict our analysis to predicting amino acid frequencies by rank rather than by amino acid identity. This result holds equally well both in alignments of empirical protein sequences and of sequences evolved under a biophysically realistic all-atom force field. Our analysis reveals a near universal shape of the frequency distributions of amino acids. This insight has the potential to lead to new models of evolution that have both increased realism and a limited number of free parameters.

## Introduction

To uncover the relationship between and the history of various protein sequences across populations and species, evolutionary biologists frequently fit mathematical models of evolution to homologous sequence alignments. Common applications of such models include phylogenetic tree reconstruction, assessment of strength and type of selection, and evolutionary rate inference. Early models had only one or two free parameters per alignment (Jukes and Cantor, 1969; Kimura, 1980), but over time models have become more complex and realistic (Goldman and Yang, 1994; Yang and Bielawski, 2000; Halpern and Bruno, 1998; Kosakovsky Pond and Frost, 2005; Yang and Nielsen, 2008; Arenas, 2015). An important insight from work in this area has been that evolving proteins display substantial variation among individual sites (Echave and Wilke, 2017), and thus site-specific models are critical. In part to address this insight, two more recent developments include mutation– selection models that estimate selection coefficients for individual amino acids at individual sites (Rodrigue et al., 2010; Rodrigue and Lartillot, 2014; Tamuri et al., 2012, 2014) and efforts to improve the biophysical realism of the models used (Koshi and Goldstein, 1998; Conant and Stadler, 2009; Meyer and Wilke, 2013; Goldstein and Pollock, 2016; Bastolla and Arenas, 2019).

One challenge with site-specific mutation–selection models is that they require the estimation of a large number of parameters, on the order of several thousand for proteins of typical lengths. Thus, they can be problematic in data-poor applications, and there is always a risk of overfitting. While this problem can be somewhat alleviated by using random-effects models (Rodrigue et al., 2010; Rodrigue and Lartillot, 2014) or penalized likelihood (Tamuri et al., 2014), it would also be useful to have simpler models that capture relevant variation at a site with only a small number of parameters. Conventionally, when simpler models are desired or needed, most researchers employ rate models that assign a single rate to each site (Pupko et al., 2002; Kosakovsky Pond and Frost, 2005; Ashkenazy et al., 2016; Spielman and Kosakovsky Pond, 2018). Rate models have been shown to provide useful summary information about evolutionary variation at individual sites, and rates can usually be recovered if selection coefficients are known (Spielman and Wilke, 2015, 2016). However, the reverse inference is generally not possible. The rate at which a site evolves does not contain any information about the amino acid distribution and/or selection coefficients at that site.

Here, we explore whether there is some avenue to capturing site variation with only one or two parameters per site while also retaining meaningful information about the amino acid distribution. To this end, we evaluate a novel approach for characterizing site-specific amino acid variation, which has previously been used to describe amino acid frequency distributions averaged across sites with similar relative solvent accessibility (RSA) (Ramsey et al., 2011). We demonstrate that a simple Boltzmann-like distribution with a single free parameter can accurately represent observed amino acid frequencies, as long as we allow for one important simplification of the problem: We rank amino acids from most abundant to least abundant at each site, and then describe the frequency distribution of the ranks, rather than of specific amino acids. We find that this approach works both for empirically collected multiple sequence alignments and for alignments generated by evolutionary simulation using a biophysically realistic, all-atom model of protein stability. We further find that introducing additional parameters into the distribution does not seem to lead to further improvements over the one-parameter description. In summary, we uncover a property of amino acid distributions that, if incorporated into models of protein evolution, could increase the realism of these models while keeping the number of free parameters limited.

## Theory

To evaluate the evolutionary and biophysical constraints acting on a site in a protein, we can assemble a large alignment of homologous sequences for that protein and then inspect the distribution of amino acids present at the site. Absent any constraint, all 20 amino acids should be present in the distribution. In reality, however, sites face various constraints, and consequently a smaller number of amino acids is actually observed (Echave and Wilke, 2017). Empirical alignments have shown that, on average, sites tend to have 3–5 amino acids that comprise the majority of the distribution. While this is true on average, the actual number of amino acids observed can vary widely and will depend on the site’s location within the protein structure.

As a simple measure of the amino acid variability at a site, we can use the effective number of amino acids *n*_eff_, defined as (Strait and Dewey, 1996; Goldstein and Pollock, 2016; Echave and Wilke, 2017)

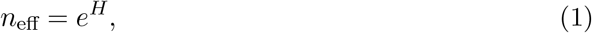

where *H* is the site entropy,

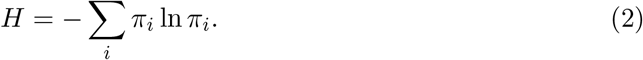

Here, the sum runs over all 20 amino acids *i*, and *π*_*i*_ represents the frequency of amino acid *i* at the focal site in the alignment. By definition, *n*_eff_ is a number between 1 and 20. A value of 1 indicates that only a single amino acid is present at the given site, and a value of 20 indicates that all 20 amino acids are present at equal frequencies. Intermediate numbers represent cases between these two extremes. For example, *n*_eff_ = 3 would indicate that the majority of the distribution is made up of three distinct amino acids, even if a fourth or a fifth amino acid may be observed at the site in very low frequencies.

The measures of site entropy and *n*_eff_ can tell us biologically relevant information about a site, such as the amount of evolutionary constraint on the site. Sites with a high *n*_eff_ can accept a wide range of different amino acids and are likely under weak selection. By contrast, sites with a low *n*_eff_ can accept only a few amino acids and are likely under strong purifying selection.

In principle, the amino acid frequencies *π*_*i*_ in Equation 2 can take on any arbitrary value. However, in practice, more structure can be assumed. First, we can write the frequencies in the form of a Boltzmann distribution with suitably defined energy levels *E*_*i*_:

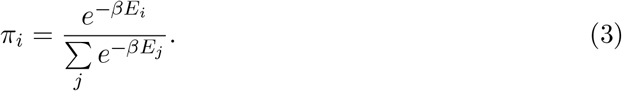

Here, *β* is the inverse temperature, and in the following we set *β* = 1 without loss of generality. Equation 3 is a mathematical identity, i.e., for any set of (non-zero) *π*_*i*_ we can define a corresponding set of *E*_*i*_ such that Equation 3 holds.

Prior work has suggested that if we order the *π*_*i*_ from largest to smallest, such that *π*_1_ *≥ π*_2_ *≥··· ≥ π*_20_, then the corresponding energy levels are approximately evenly spaced, such that *E*_*i*_ = *iλ* with a site-specific numerical constant λ (Porto et al., 2005; Ramsey et al., 2011). Ramsey et al. (2011) found this description to work well when amino acid distributions were averaged over many sites with comparable solvent accessibility. We assume here the result holds similarly for site-specific frequencies. Thus, we rewrite Equation 3 as

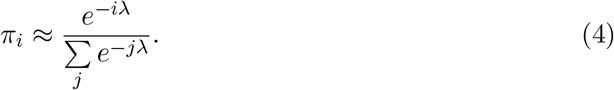

Larger values of λ correspond to sites with a smaller effective number of amino acids present, i.e., more conserved sites. There is a systematic trend of λ to increase as we move from the surface of a protein to its core (Ramsey et al., 2011).

Equation 4 predicts the amino acid distribution at a site from a single free parameter, λ. Consequently, we can calculate the *n*_eff_ that corresponds to a given λ. By substituting Equation 4 into Equation 2 and then 1, we obtain

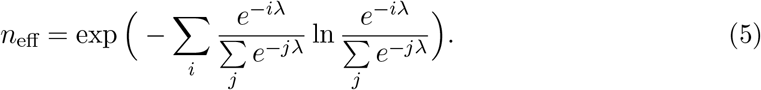

This equation is exact, i.e., Equation 5 provides us with the exact *n*_eff_ corresponding to a given λ, assuming Equation 4 is true. However, Equation 5 also suggests that the inverse relationship may be true as well. Given an *n*_eff_, which we can calculate from a column in a multiple sequence alignment, we can invert Equation 5 to predict a corresponding λ. This inversion cannot be done analytically, but it is straightforward numerically.

## Results

The preceding section suggests that the amino acid distribution at a site can be represented by a single parameter λ, where once we know λ we know the individual frequencies *π*_*i*_, modulo a reordering. (I.e., we will not know which amino acid is the most frequent or the second-most frequent etc.; we will only know what their relative frequencies are.) Note that this is a stronger statement than saying we can represent a site by its *n*_eff_, because while *n*_eff_ captures the variability at a site it does not a priori fix the frequencies *π*_*i*_.

We can test this prediction empirically by taking columns in multiple sequence alignments (MSAs), calculating *n*_eff_ values, converting them into λ values, then calculating predicted amino acid frequencies from the λ values and assessing how close they are to the observed frequencies. For this purpose, we here use previously published MSAs of taxonomically and functionally diverse proteins (Jiang et al., 2018). Specifically, we analyze MSAs corresponding to 10 arbitrarily chosen proteins (Table 1), and we consider both MSAs of natural sequences and of sequences generated via simulation, using an accelerated origin-fixation algorithm (Teufel and Wilke, 2017). Although the natural sequences are subjected to mutational biases and codon degeneracy and the simulated sequences are not (all mutations between amino acids are equally likely), only minor differences in amino acid frequencies have been observed between these sets of alignments (Jiang et al., 2018). For this reason, we take observed amino acid frequencies as is and do not correct for mutation biases.

**Table 1:**
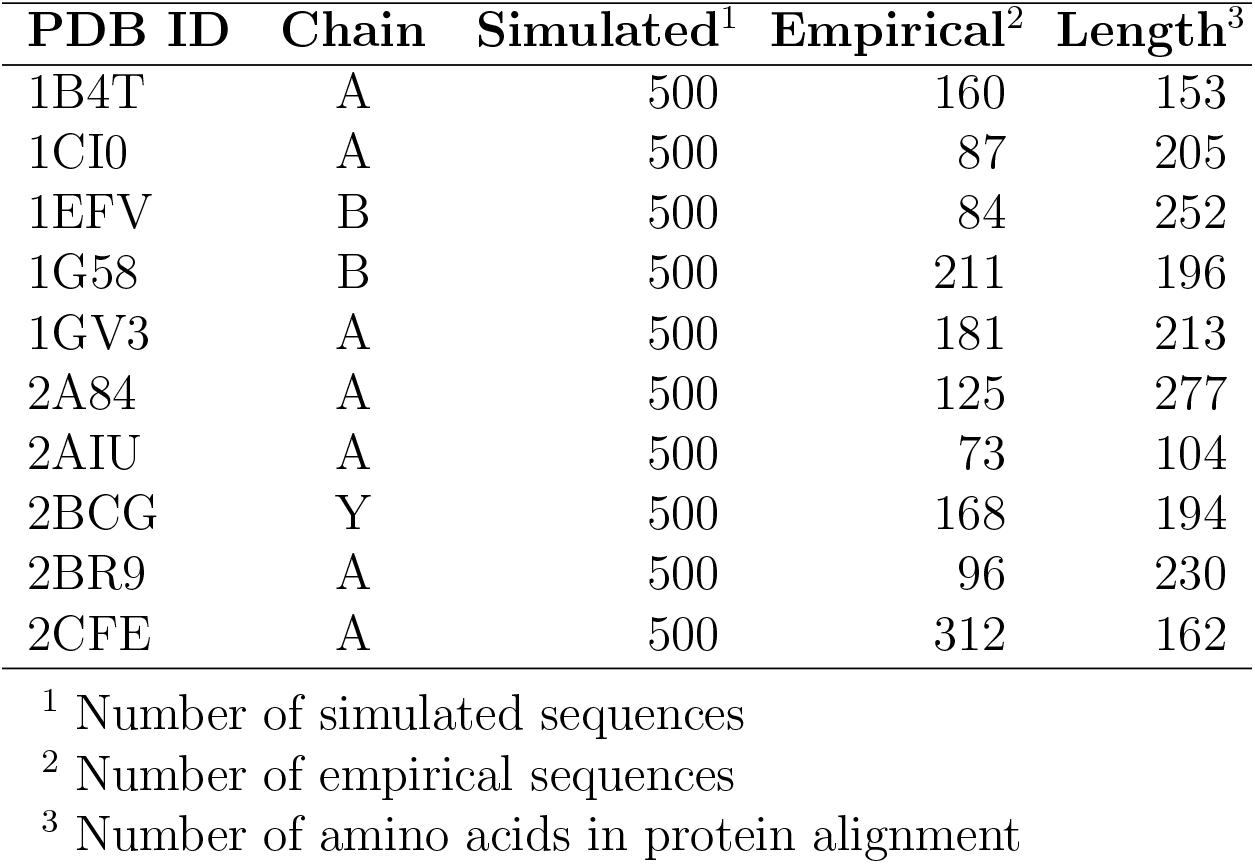
Identity and characteristics of the pairs of simulated and empirical protein alignments analyzed here. Data from Jiang et al. (2018).

### Theoretical expectation observed in simulated and empirical alignments

Because alignments of natural sequences are confounded by phylogenetic relatedness and the simulated alignments are not (they were created using a star phylogeny), we expect that our approach will fit the simulated alignments better than the natural ones. Therefore, we first apply it to simulated sequences.

We begin with simulated sequences corresponding to a yeast copper-zinc superoxide dismutase (PDB ID: 1B4T). The MSA has 153 sites, and we observe substantial heterogeneity among the sites in terms of the number and type of amino acids present. To demonstrate our theoretical reasoning described in the preceding section, we first consider one of the variable sites (site 35), perform a linear regression on the ranked, log-transformed frequencies to obtain λ, and then compare the distribution given by Equation 4 to the empirical distribution at the site (Fig. 1). The two distributions look visually similar, and a *χ*^2^ goodness-of-fit test detects no significant difference between them (*p* = 0.106).

**Figure 1:**
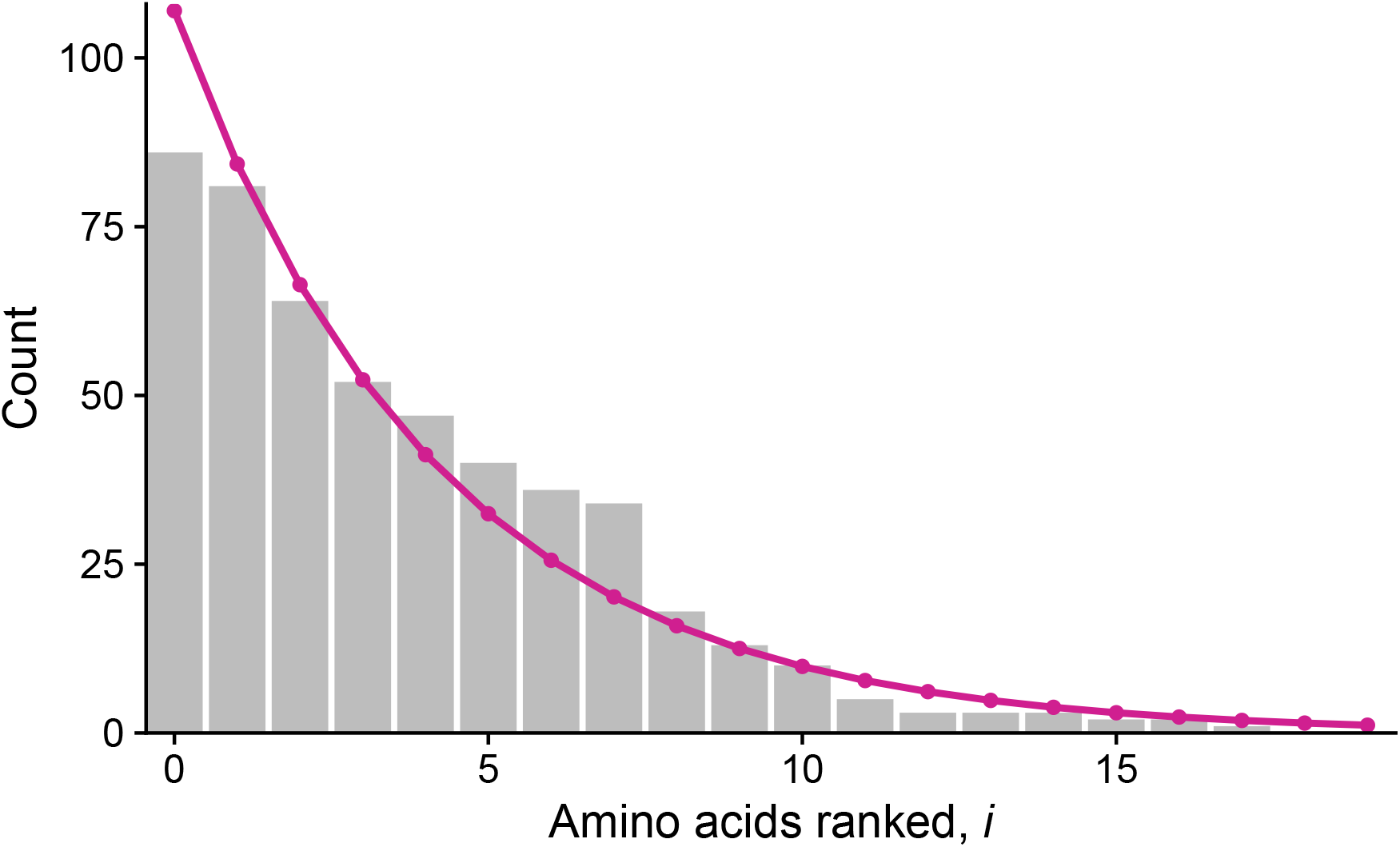
Site-specific distribution fit to observed amino acid counts. The gray bars represent the counts observed in 500 simulated sequences, while the magenta line depicts the distribution approximated from the linear regression fit to the log-transformed count data. The plot above is for site 35 for the simulated alignment of a yeast copper-zinc superoxide dismutase (PDB ID: 1B4T).

Next, we repeat this procedure at all remaining sites in this protein’s alignment. To visualize the results from this analysis, we take the λ values obtained from the linear regressions and plot 1*/*λ against the *n*_eff_ calculated directly from the observed amino acid frequencies (Fig. 2). We plot 1*/*λ instead of λ to avoid the divergence at *n*_eff_ = 1, where 1*/*λ = 0. We find that the majority of sites fall near the line defined by Equation 5, which is expected if the theoretical relationship between *n*_eff_ and λ holds true (Fig. 2). However, even though all sites are near this theoretical relationship, there are measurable deviations from the exact theory. Many sites fail the *χ*^2^ goodness-of-fit test after False-Discovery-Rate (FDR) correction, indicated in orange color (Fig. 2). As a general pattern, we observe that sites with a higher effective number of amino acids are more likely to not fail the *χ*^2^ test (blue points).

**Figure 2:**
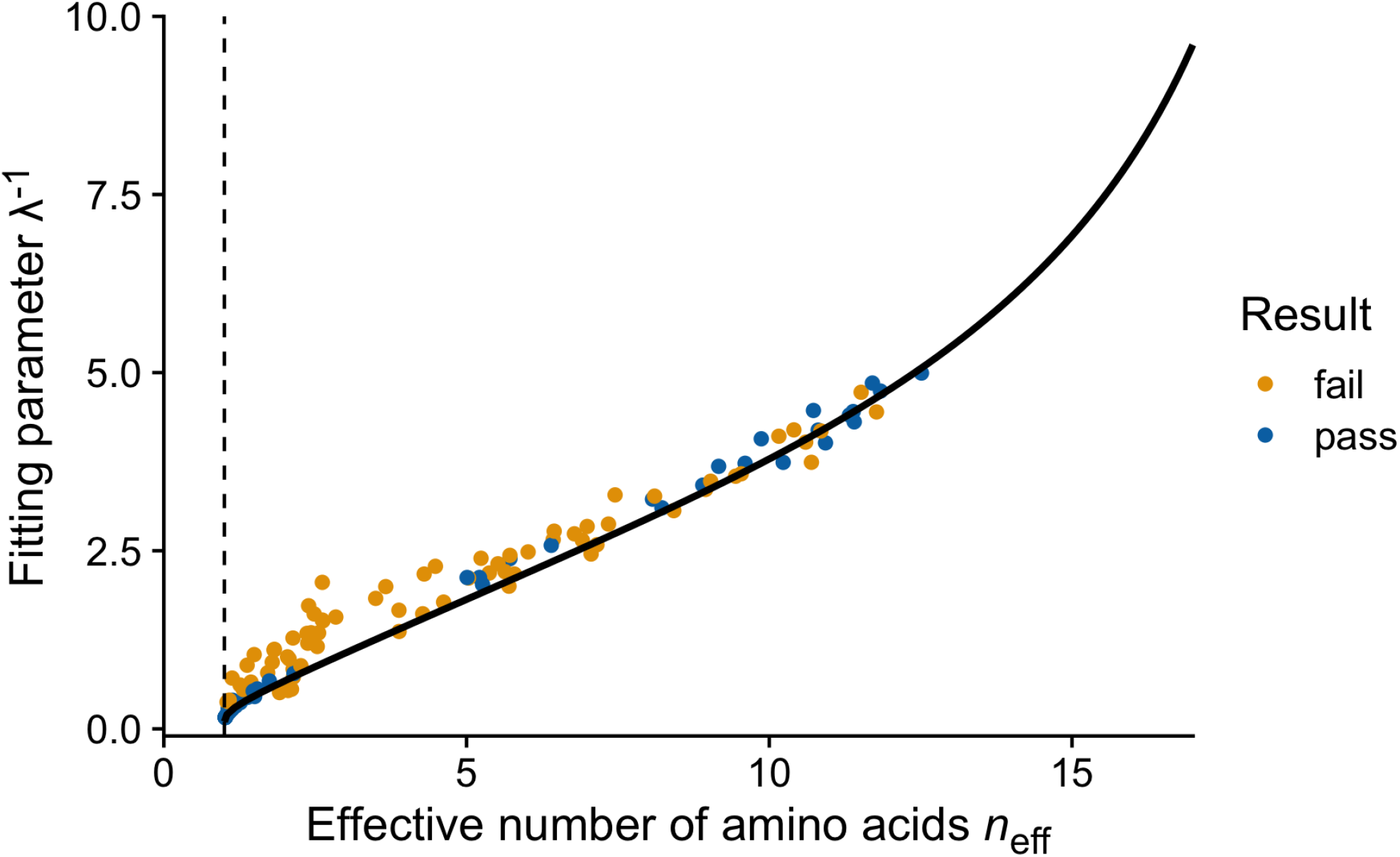
Relationship between the effective number of amino acids (*n*_eff_) and λ^−1^, for the simulated alignment from Figure 1 (PDB ID: 1B4T). Each point represents a site in the alignment with λ values from a linear regression on the transformed count data and *n*_eff_ values calculated with Equation 1 on observed counts. The use of λ^−1^ improves our ability to visualize this relationship for small values of *n*_eff_ where λ diverges. The black line represents the theoretical expectation calculated from Equation 5. Orange points indicate sites that fail the *χ*^2^ test (75 sites), while blue sites show no significant difference between the actual and the fit distribution (50 sites). Sites with only one amino acid present were excluded from the *χ*^2^ analysis (29 sites).

We find similar results in nine additional simulated protein alignments (Fig. 3). Most sites fall close to the line defined by Equation 5, but nevertheless, only a moderate number of sites at high *n*_eff_ pass the *χ*^2^ test. Surprisingly, our results look better for the empirical alignments for the same ten proteins (Fig. S1). There the majority of the sites pass the *χ*^2^ test for each protein (Fig. 4 and Fig. S2). This difference is likely driven by the difference in the number of sequences in each alignment. Most empirical alignments contain between 80 and 200 sequences, whereas all simulated alignments contain 500 sequences (Table 1). We test this hypothesis by downsampling the size of each simulated alignment to match the number of sequences in its corresponding empirical alignment and repeating the analysis. We confirm that indeed the downsampled simulated alignments more closely resemble their empirical counterparts than the full simulated alignments (Fig. 4 and Fig. S2). Additionally, we find that distributions of the adjusted coefficients of determination (*R*^2^) for these regression analyses look similar across simulated and empirical alignments, with added variation in some downsampled simulated alignments (Fig. S3).

**Figure 3:**
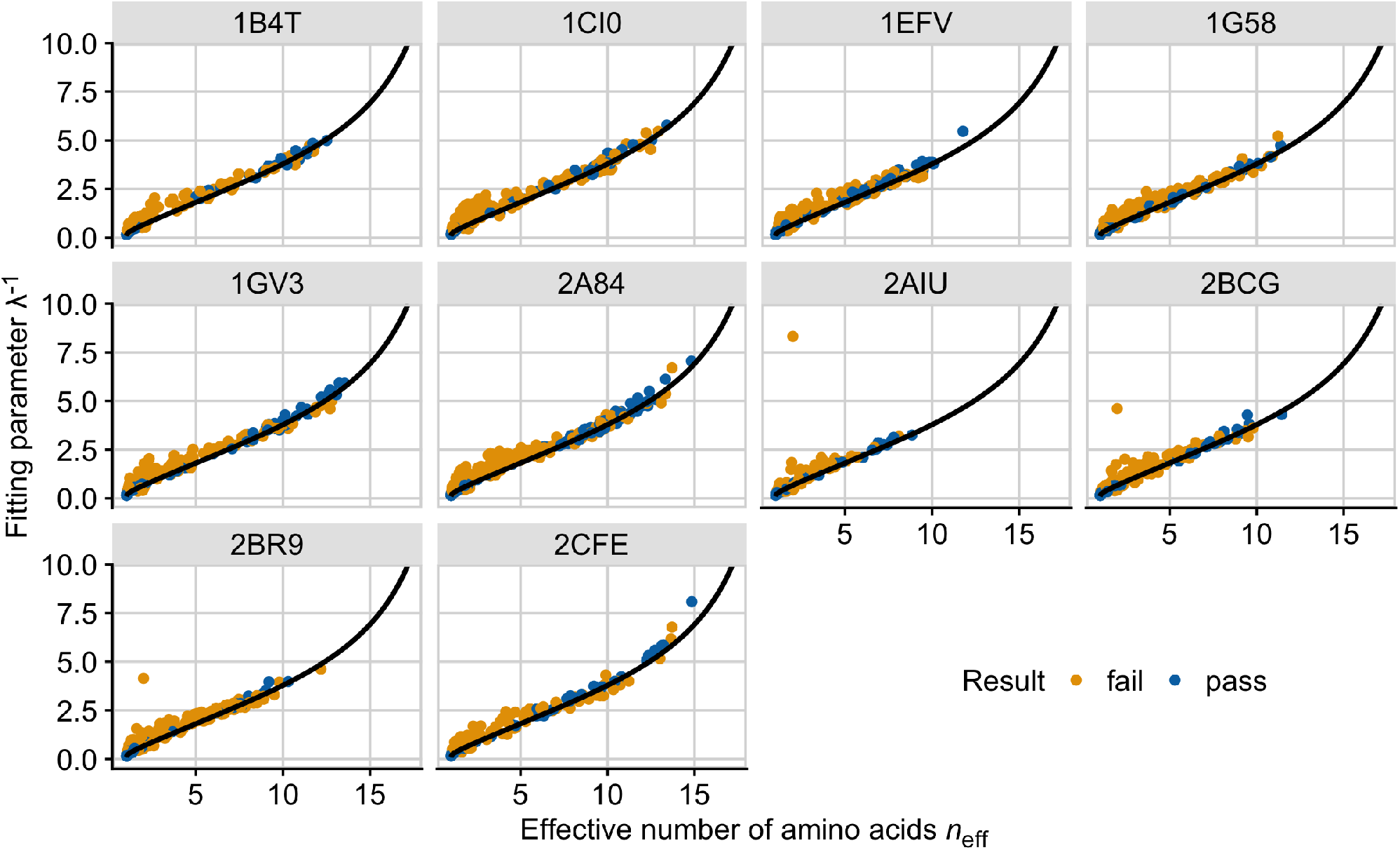
The fit of 10 simulated protein alignments to the theoretical expectation from Equation 5 (black line). The *x* axis represents the effective number of amino acids, *n*_eff_, while the *y* axis is 1*/*λ. The title of each panel indicates the protein’s identifier in the Protein Data Bank. Each point represents the distribution at a single site, where λ is the slope parameter from the linear regression and *n*_eff_ is calculated with Equation 1. Orange points indicate a failed *χ*^2^, while blue points indicate a passed *χ*^2^ after FDR correction.

**Figure 4:**
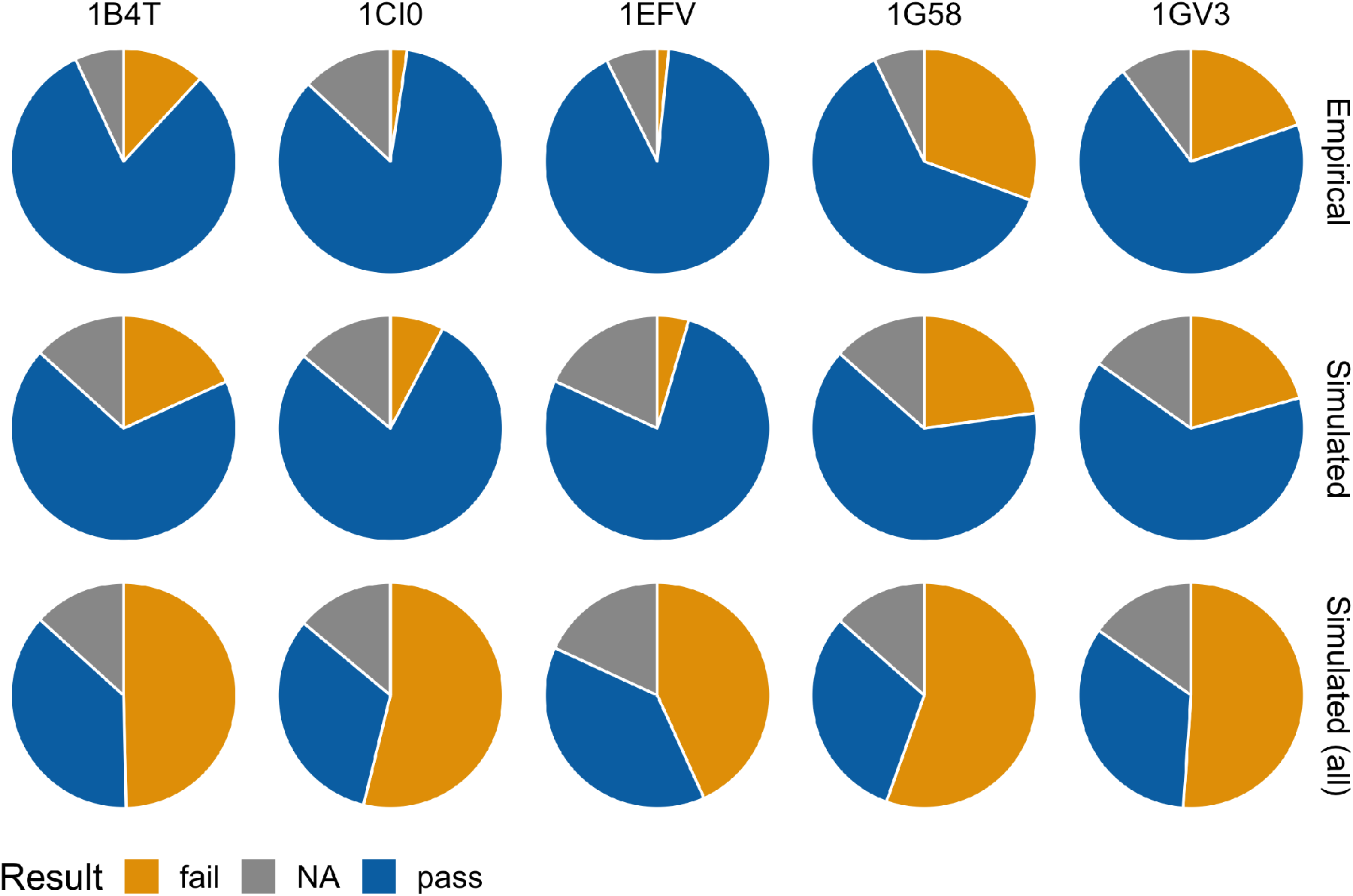
Performance of a *χ*^2^ goodness-of-fit test between the actual and estimated amino acid distributions at each site in five different proteins. Analysis was performed on actual counts observed in the alignment and the counts estimated from the linear regression. Results are compared for an empirical alignment, a simulated alignment with an equivalent number of sequences, and a simulated alignment with 500 sequences for each protein considered. A FDR correction controlled for multiple testing. Orange indicates sites that failed the *χ*^2^, while blue indicates sites that passed the *χ*^2^ and grey indicates the number of sites that could not be tested under the *χ*^2^ goodness-of-fit test due to the presence of a single amino acid.

### Null distributions confirm theoretical expectations

To test the validity of our fitting procedure, we repeat the methods discussed above under two different null distributions. As a positive control, we randomly draw amino acid distributions according to Equation 4 and then repeat our fitting procedure for these distributions. As expected, *χ*^2^ tests show no significant difference between the simulated and the fitted distributions, for all sites tested (Fig. 5, left panel).

**Figure 5:**
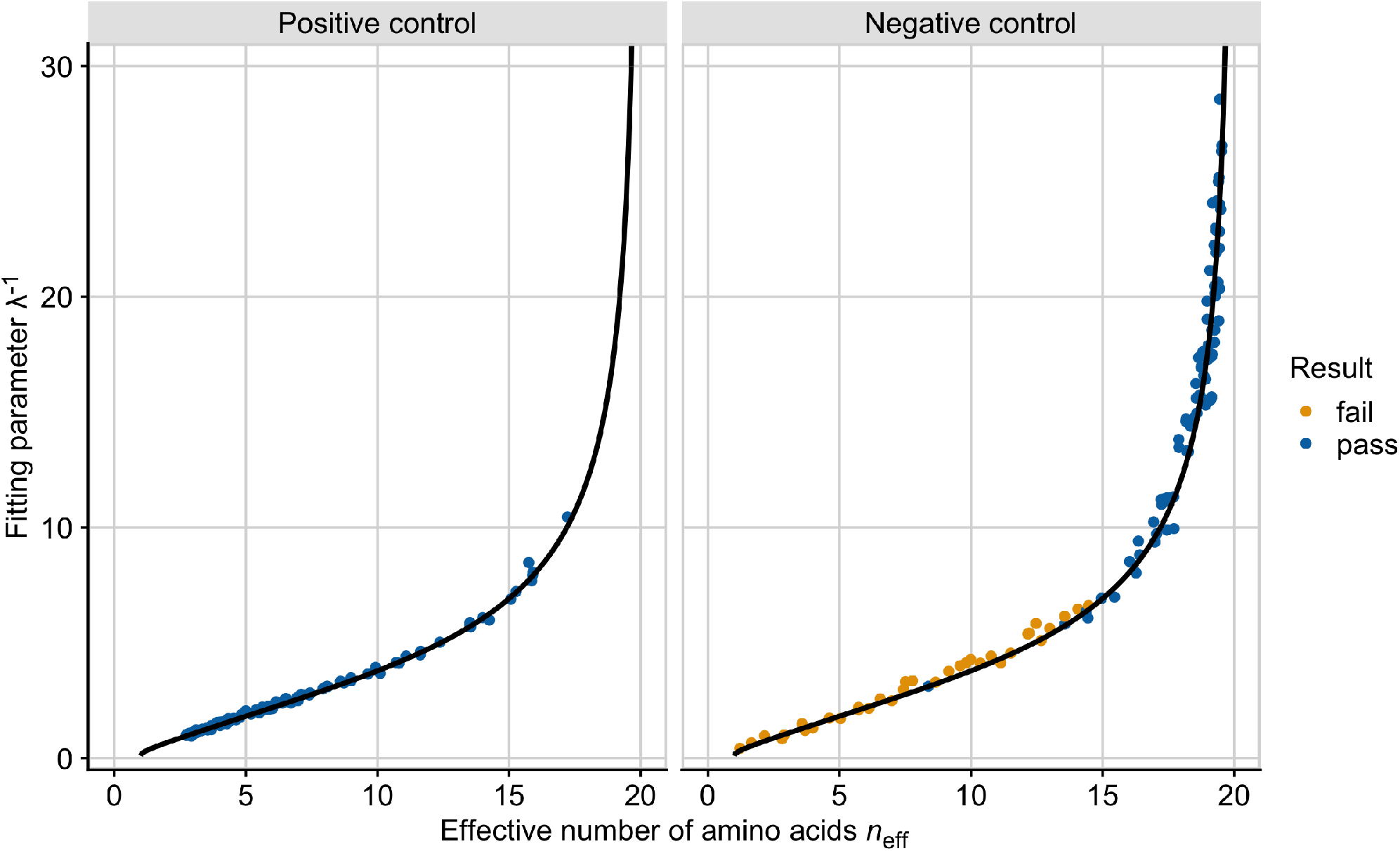
Fit of null distributions to the theoretical expectation (black line, Eq. 5). The *x* axis represents the effective number of amino acids, *n*_eff_, while the *y* axis is 1*/*λ, where λ is the slope parameter from the linear regression. Orange points indicate a failed *χ*^2^, while blue points indicate a passed *χ*^2^ after FDR correction. The plot on the left show sites with distributions simulated from Equation 4, while the distributions represented on the right were simulated from a Gaussian distribution. For each site, values of λ come from linear regression on transformed counts, while *n*_eff_ is calculated with Equation 1 on raw counts.

As a negative control, we additionally simulate amino acid counts described by a Gaussian function and then subject them to the same fitting procedure. We define expected amino acid frequencies as *π*_*i*_ ~ exp[−(*i* − 10)^2^*/σ*^2^], where *i* is an integer taking on values between 0 and 19. We use multinomial sampling to generate specific amino acid counts and then re-rank counts from largest to smallest (see Methods for details). Depending on the value of *σ* chosen, this approach can result in unrealistic amino acid distributions that are near uniform, with *n*_eff_ values of 15–20 (Fig. 5, right panel). In this previously unobserved area of parameter space, we find that the simulated distributions are accurately represented by our theoretical relationship between 1*/*λ and *n*_eff_. This is the case because for near-uniform amino acid distributions, the quadratic term in the Gaussian can be neglected. By contrast, for parameter choices that result in smaller *n*_eff_ values, we now see consistent deviations between the observed and the expected distributions (most sites fail the *χ*^2^ test and are colored in orange), even though all sites fall near the expected relationship between 1*/*λ and *n*_eff_ (Fig. 5, right panel). This result demonstrates that our *χ*^2^ test is sensitive to subtle deviations in amino acid frequencies from the distribution given by Equation 4.

### Additional parameters and non-linearity do not improve regression fit

As an additional method of assessing how well our characterization describes actual amino acid distributions, we can also compare the *n*_eff_ values calculated from an empirical alignment to the *n*_eff_ values calculated from the frequencies of the fitted distribution. If the fitted distribution accurately reflects the empirical distribution, then these two sets of *n*_eff_ values should be very similar to each other. In fact, we generally find that they are highly correlated and near the *x* = *y* line (e.g., *R*^2^ = 0.971, Fig. 6 top panel for the MSA corresponding to yeast copper-zinc superoxide dismutase, PDB ID: 1B4T).

**Figure 6:**
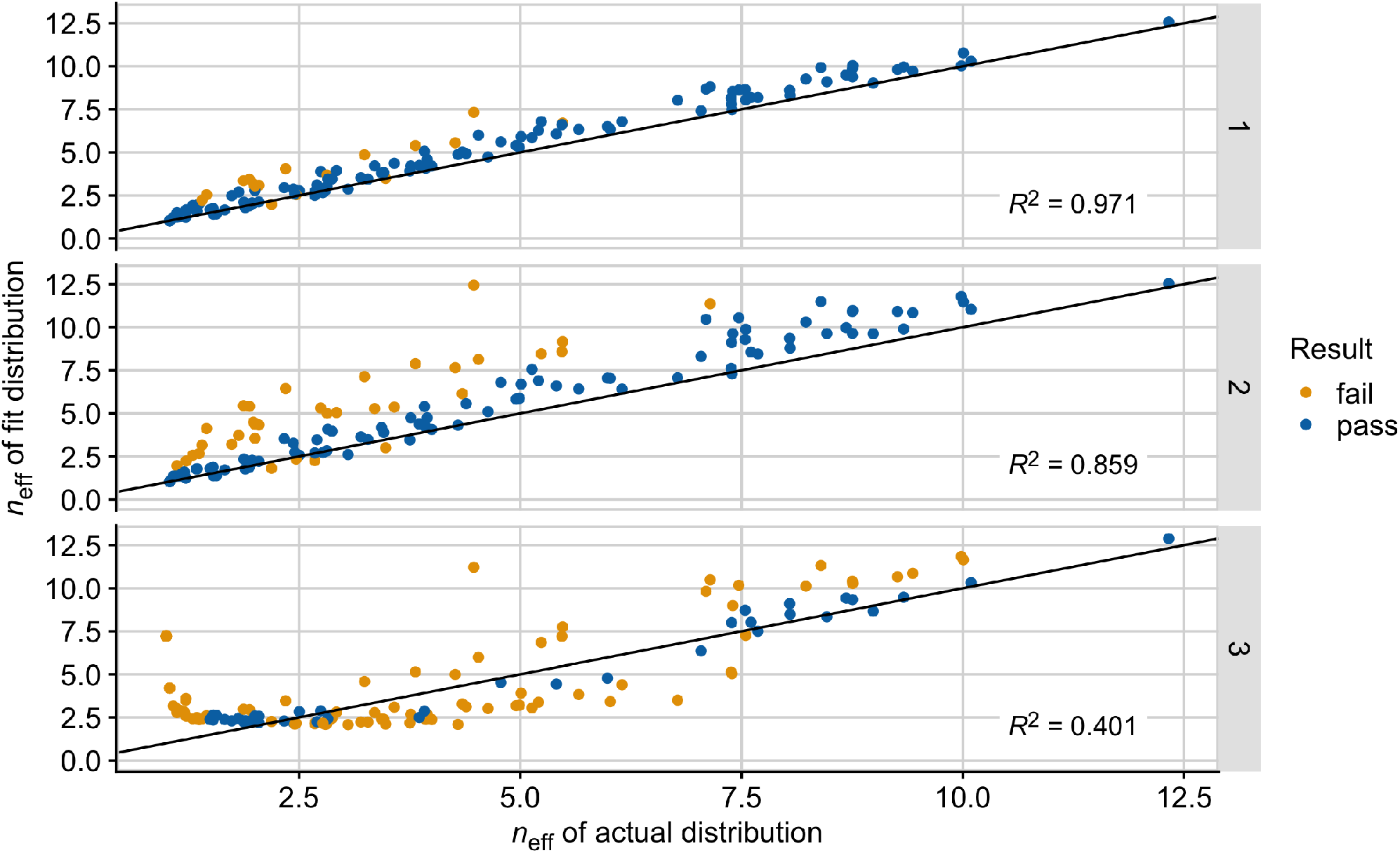
Conservation of the effective number of amino acids (*n*_eff_) in the fit distribution. For each site, *n*_eff_ was calculated with Equation 1 on the observed counts in the alignment and on the counts estimated from the linear regression. Both measures are presented for each site in the empirical 1B4T alignment when the distribution was fit with either one, two, or three parameters, as indicated by the labels on the right. The black lines represent the points for which *x* = *y*, and the *R*^2^ reported in the figure is from the correlation between *n*_eff_ in the actual and fit distribution. Colors indicate the result of the *χ*^2^ goodness-of-fit test with FDR correction.

While the high correlations we observe are an excellent result, we can ask if we could do better. For the top panel in Fig. 6, of the 153 sites in the alignment 20 had to be excluded from the comparison as they are completely conserved for a single amino acid and the fitting procedure fails in this case. Of the remaining 133 sites, 17 sites (12.8%) fail the *χ*^2^ test. Notably, the fits for this result are done using a one-parameter model, where normalized, log-transformed counts are fit to a model without intercept, 0 − λ*i* (*i* is an integer running from 0 to 19, see Methods for details). A single parameter fit tends to performs poorly on sites that are highly conserved for either one or two equally frequent amino acids.

To explore alternatives, we can make two modifications to this analysis: First, we can perform a different linear regression that is however still consistent with Equation 4, a two-parameter model with an intercept term: *b* − λ*i*. The intercept term cancels out in Equation 4, but it affects the λ value and thus the amino acid frequencies obtained during fitting. Second, we can test a quadratic regression where a third parameter, *c*, is added to capture the shape of the curve: *b* − λ*i* + *ci*^2^.

We find that both the two-parameter regression and the three-parameter regression perform worse than the one-parameter regression based on several metrics: *p*-values from the *χ*^2^ goodness-of-fit test, *R*^2^ values on transformed count data from regression analyses, and *R*^2^ values from the correlation between actual and estimated *n*_eff_ values (Fig. S4 and 6). For the two-parameter regression, we are still unable to fit the 20 sites with only one amino acid observed. Of the remaining 133 sites, 34 sites (25.6%) fail the *χ*^2^ test, and the *R*^2^ between the *n*_eff_ calculated on the actual and fit distribution has declined to *R*^2^ = 0.859. The three-parameter regression leads to the worst performance overall: 110 of 153 sites (71.9%) fail the *χ*^2^ test, and the *R*^2^ between the *n*_eff_ calculated on the actual and fit distribution has declined to *R*^2^ = 0.401.

As a general pattern, we observe that additional parameters and non-linearity in the fitted model lead to increased variability in our ability to recapture the *n*_eff_ of the actual distribution. With one parameter, we tend to slightly overestimate *n*_eff_ with our fitted distribution, in particular towards low *n*_eff_ values (Fig. 6, top). The addition of the intercept parameter moves points further from the *x* = *y* line overall (Fig. 6, middle). Finally, the quadratic regression, with three parameters, produces additional spread in both directions, resulting in quite severe over- and underestimations of *n*_eff_. We find that these results hold true across all 10 empirical proteins considered (Fig. S5). A small constant was added to amino acid counts prior to fitting only in the case of our three-parameter regression as it was necessary to obtain enough data points for fitting at each site (*n* > 2). This constant was excluded from the one- and two-parameter regressions as it was not required for fitting and was shown to decrease our ability to fit the distribution (Fig. S6, top and middle panel). We additionally attempted to improve our three-parameter fit through regularization with ridge regression, which however increased the upward bias in the *n*_eff_ of the fitted distribution (Fig. S6, bottom panel) (Friedman et al., 2010). Thus, in conclusion, a simple one-parameter linear regression without intercept maximizes our ability to capture the empirical amino acid distribution.

## Discussion

We have shown that a simple Boltzmann-like distribution with a single free parameter works surprisingly well at capturing amino acid variability at individual sites in protein multiple sequence alignments, as long as we are prepared to ignore amino acid identity and simply order amino acids from most frequent to least frequent. We have found that this description works both in empirical alignments and in alignments derived from simulations using a physics-based atom-level model of protein structure. For many, though not all, sites in an alignment, the single one-parameter description is sufficiently accurate to be statistically indistinguishable from the true amino acid distribution. In general, deviations from the true distribution tend to be more prominent at sites that display less variability overall.

That amino acid frequencies should be Boltzmann distributed follows from theoretical models of protein stability (Dokholyan and Shakhnovich, 2001; Dokholyan et al., 2002; Echave et al., 2015). However, we note that our finding here is stricter than prior theoretical predictions. In the most general scenario, we can write any arbitrary amino acid distribution in Boltzmann form, by appropriately choosing energy levels for each individual amino acid (see Eq. 3). Here, instead, we are arguing that energy levels are approximately uniformly spaced, so that a single parameter λ (corresponding, in effect, to the distance between two energy levels) can capture the entire distribution at a site. Previous work has shown that energy levels are approximately evenly spaced for amino acid distributions averaged across sites with similar relative solvent accessibility (Ramsey et al., 2011). We find that this observation also holds true for site-specific amino acid distributions in both simulated and empirical protein sequences. The mechanisms driving this empirical observation remain unknown and should be explored in future work.

While our approach is able to reproduce the general shape of the distribution of amino acids at a site, the actual and the estimated distribution are not always an exact match. There are subtle deviations from the simple exponential decay that can be detected when alignments are sufficiently large. In particular, while the majority of sites in simulated and empirical alignments pass a *χ*^2^ goodness-of-fit test when a limited number of sequences are used in the regression (fewer than *~* 200), a larger fraction of sites fails the *χ*^2^ goodness-offit test for simulated alignments with the full 500 sequences. We have further found that the distribution of amino acids is best characterized when fit (after log-transform) with a simple linear regression without intercept. Adding a second parameter (the intercept) to the linear regression or implementing a quadratic (three parameter) regression to aid in approximating the distribution can improve *χ*^2^ results for some sites, in particular highly conserved sites. However, on average, the two- and three-parameter regressions perform worse than the one-parameter regression.

The characterization presented here allows for a general approximation of the shape of amino acid distributions. However, these theoretical distributions are not exact and vary in their ability to recover the observed amino acid frequencies in an empirical or simulated alignment. The biggest challenges seem to arise at sites that are highly conserved for only a few amino acids. Unfortunately, such sites are common in empirical alignments. In addition, just like is the case with evolutionary rate measures, approximating an amino acid distribution by rank does not retain any information about the identity of amino acids found at individual sites. While predicting the rank order of amino acids is beyond the scope of this research, it might be possible to develop standard rankings based on the biology of the system. For example, we observe that amino acids occupy rank 0 at different frequencies based on their location within the protein structure (Fig. S7), e.g., hydrophobic amino acids tend to be most abundant at sites that are buried in the protein core. Potential strategies for jointly predicting rank frequencies and identities in evolutionary models are discussed below.

The ability to reduce an amino acid distribution from 20 parameters to one might be useful if applied to models of protein evolution. Current phenomenological models of protein evolution that require only a few parameters, such as *dN/dS* models (Goldman and Yang, 1994; Kimura, 1977; Yang and Bielawski, 2000; Kryazhimskiy and Plotkin, 2008), provide aggregate information about how sites have changed but capture little information about the distribution of amino acids at individual sites (Arenas and Posada, 2014; Halpern and Bruno, 1998). Additionally, they tend to be sensitive to evolutionary forces if not explicitly modeled (Wilson and McVean, 2006; Arenas, 2015; Spielman and Wilke, 2015; Kryazhimskiy and Plotkin, 2008). Alternatively, mechanistic mutation–selection models often rely on numerous parameters to account for observed heterogeneity in amino acid frequencies across sites (Halpern and Bruno, 1998; Rodrigue et al., 2010; Yang and Nielsen, 2008; Bruno, 1996; Tamuri et al., 2012, 2014). These models connect amino acid frequencies with fitness, but they are computationally expensive and can result in an over-parameterized representation of the sequence space (Puller et al., 2020; Rodrigue, 2013; Spielman and Wilke, 2016). We note that a site-specific model with one free parameter has previously been developed for phylogenetic inference (Arenas et al., 2015). By accounting for structural properties, this mean-field model improves inference of evolutionary events, but its estimates of site-specific sequence entropy and substitution rate disagree with empirical data (Jimenez et al., 2018).

Current approaches to preventing over-parameterization of mutation–selection models include using random-effects models (which in effect share amino acid distributions among sites, Rodrigue et al. 2010) and regularization (which causes low amino acid frequencies to be set to zero, Tamuri et al. 2014). Both approaches work reasonably well, but random-effects models have a tendency to produce too many non-zero amino acid frequencies (Spielman and Wilke, 2016). Our results here suggest a mathematical explanation for this observation: Random-effects models tend to represent amino acid distributions as a weighted average over a finite set of propensity vectors as the true propensity vector is unknown (Rodrigue et al., 2010). While individual propensity vectors can converge into a Boltzmann form, a weighted average of different Boltzmann distributions—with amino acids in different rank order—will generally not be Boltzmann, and in fact will over-populate rare amino acids.

We emphasize that we have not proposed a new model of evolution here. We have merely identified a property of empirical amino acid distributions that models of evolution should be able to capture. Integrating this characterization into such models could improve our ability to infer phylogenetic relationships as well as evolutionary rates and processes. We would like to suggest some avenues of how our insights could be incorporated into future models. First, for random-effects models, instead of using arbitrary propensity vectors it might be useful to enforce the Boltzmann form with a single free parameter λ, and then fit the amino acid order at each site. Amino acid order can be represented by permutations, and permutations can be sampled using standard MCMC approaches (Strauß et al., 2019). Of course, any models that fit amino acid order would require more than one parameter per site.

For fixed-effects models, on the other hand, it may be possible to derive a new regularization approach that uses our Eq. 4 for regularizing amino acid frequencies. We note that fixed-effects models for phylogenetic inference often lack the statistical guarantees of traditional likelihood estimation and thus may be prone to overfitting based on the dimensionality of the site-specific variables (Rodrigue, 2013). Researchers wishing to incorporate our characterization into such models should do so with caution.

## Methods

### Origin of multiple sequence alignments

We use protein multiple sequence alignments (MSAs) from an existing data set that contains both empirical and simulated alignments for the same protein structures (Ramsey et al., 2011; Jiang et al., 2018). The empirical alignments were originally assembled by Ramsey et al. (2011) and include 38 MSAs, each containing at least 50 sequences, for 38 distinct protein structures (Jackson et al., 2013). For each protein structure, Jiang et al. (2018) subsequently added a simulated alignment containing 500 sequences. The simulated alignments were simulated using an accelerated origin-fixation algorithm (Jiang et al., 2018; Teufel and Wilke, 2017) with a physics-based, atom-level model of protein structure as a fitness function. The simulations were performed along a star phylogeny, such that no phylogenetic structure is present in the simulated alignments (Jiang et al., 2018). The empirical alignments had originally been filtered to remove sequences of ≥ 80% similarity, so phylogenetic structure in these alignments is limited but not absent (Ramsey et al., 2011).

Here, we arbitrarily selected 10 protein structures with their associated empirical and simulated alignments for further analysis (Table 1). We note that because empirical alignments contain gaps, the total number of amino acids observed at any given site might be smaller than the number sequences listed.

### Approximating the distribution of amino acids at a site

We use the following fitting procedure to estimate the site-specific constant λ that parameterizes our Boltzmann-like amino acid distribution (Equation 4). Rather than fitting an exponential model directly to the observed count data, we transform the data as described below for use in a linear regression. At each site in an alignment, we first count all amino acids and then rank them by frequency, from most to least abundant. These ranked counts are then rescaled relative to the most abundant amino acid at that site and subsequently log-transformed. We then fit a linear regression of the form 0 − λ*i* for *i* = 0, 1,…, 19 to the rescaled and transformed counts, excluding any amino acids or sequences (gaps) that were not observed in the alignment. The 0 in front of −λ*i* indicates the absence of an intercept term, i.e., the regression model is forced to go through 0 at *i* = 0.

Once we have obtained a λ value at a site, we calculate expected amino acid counts by computing −*λe*^−λ*i*^ for *i* = 0, 1,…, 19. For direct comparison with observed counts, the obtained values need to be normalized relative to their sum and multiplied by the total number of counts at that site in the observed data (this is equivalent to the number of sequences in simulated MSAs, but varies based on the presence of gaps in empirical MSAs). Because we are fitting with realized frequencies instead of equilibrium frequencies, the λ values calculated here are likely subject to some degree of sampling bias.

We assess the goodness-of-fit of the fitted to the observed distribution at each site by comparing expected with observed amino acid counts using a *χ*^2^ test with 18 degrees of freedom. We correct for multiple testing via the False-Discovery-Rate (FDR) correction (Benjamini and Hochberg, 1995). A *χ*^2^ test on count data is the appropriate test here as we are interested in our ability to reproduce the original distribution from λ alone. We additionally report the proportion of variance explained (*R*^2^) from the linear regression on the observed and expected log-transformed counts.

### Generating null distributions

To generate null distributions, we specify amino acid frequencies *π*_*i*_ at each site and then draw amino acid counts for *n* = 500 sequences from multinomial distributions with expected counts *nπ*_*i*_. In all null distributions, we use 110 distinct sites. After generating the samples under various distributions, we follow the same procedures for our empirical and simulated alignments: We transform the data by ranking counts from most to least frequent, normalizing, and log-transforming prior to fitting.

As a positive control, we use *π*_*i*_ = *Ce*^−λ*i*^, with *C* chosen such that Σ_*i*_ *π*_*i*_ = 1. We generate 110 values of λ uniformly spaced from 0.1 to 1. This choice guarantees that the resulting distribution of *n*_eff_ looks similar to those observed in our protein alignments. Each value of λ is used to simulate a site within a 500 sequence alignment.

As a negative control, we use 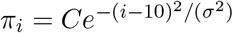. The constant *C* is again chosen such that ∑_*i*_ *π*_*i*_ = 1. For small *σ*, this choice generates amino acid frequencies that decay faster than exponential. For large *σ*, on the other hand, this choice generates frequencies that are nearly uniform. We select 110 *σ* values uniformly spaced from 0.5 to 15 to simulate sites with values of *n*_eff_ ranging from 1 to 19.

### Varying the number of parameters in fitting

We modify the fitting procedures described above by including additional parameters and non-linearity during our analysis on empirical sequence alignments. Under our normal procedure, we fit 0 *–λi* to the observed amino acid count data, as explained above. We can incorporate a second parameter into this linear regression by relaxing the assumption of a zero intercept term. Thus, we fit *b − λi* to the ranked, normalized, and log-transformed counts for only those amino acids that were observed in our data. We obtain expected counts from this fit by computing *e*^*b*–λ*i*^ for amino acids *i* = 0, 1,…, 19, normalizing to their relative frequency, and rescaling with the total number of count observations for each site.

To capture non-linear effects in the distributions, we additionally fit a three-parameter quadratic regression: *b* – λ*i* + *ci*^2^. While the previous regressions could be implemented on a small number of observed amino acids, a quadratic fit requires more data points (i.e., more non-zero amino acid counts). To deal with this limitation, we add a small constant (0.01) to all amino acid counts at all sites prior to data transformation and fitting. With this addition, the updated count values are ranked, normalized, and log-transformed prior to fitting, resulting in all 20 amino acids being present (and fit) at all sites. After obtaining λ, *b*, and *c* from the fit at each site, we compute 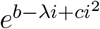 for *i* = 0, 1,…, 19, normalize each count based on its relative frequency, and rescale with the total number of counts observed at that site.

The results of all three fitting procedures are evaluated on their performance approximating the observed distribution via *χ*^2^ goodness-of-fit tests. We test our ability to recover the effective number of amino acids by calculating *n*_eff_ on both the actual and estimated distribution for each site as described by Equation 1. The relationship between these measures at each site is compared across fitting methods and presented with their respective coefficients of determination for all empirical alignments.

Results of Figure S6 are obtained by re-running the one- and two-parameter regressions on amino acid counts after a small constant (0.01) was added (top and middle panel, respectively). As in the three-parameter case, this results in all 20 amino acids being fit at each site. We additionally modify the three-parameter regression described above by fitting *b* – λ*i* + *ci*^2^ with ridge regression in the glmnet R package (Friedman et al., 2010). In order to not add an additional parameter to our analysis, we perform *k*-fold cross validation to identify the optimal tuning parameter (λ_tuning_) at each site with more than one amino acid observed, and select one value for use across all sites (Fig. S8). Based on the observed distribution, we fit the ridge regression in glmnet with λ_tuning_ = 0.28 and *α* = 0 at each site (Fig. S6, bottom panel). The expected number of amino acids at each site is calculated from the values of *b*, λ, and *c* obtained from the ridge regression.

## Code

Data and code for this work are available at: https://github.com/mmjohn/amino-acid-distributions. All data analysis and figure production was performed in R (R Core Team, 2019), making extensive use of the tidyverse family of packages (Wickham et al., 2019).

## Acknowledgements

This work was supported by National Institutes of Health (NIH) grant R01 GM088344. M.M.J. acknowledges support from NIH training grant T32 LM012414-01A1.

## Supplemental Figures

**Figure S1:**
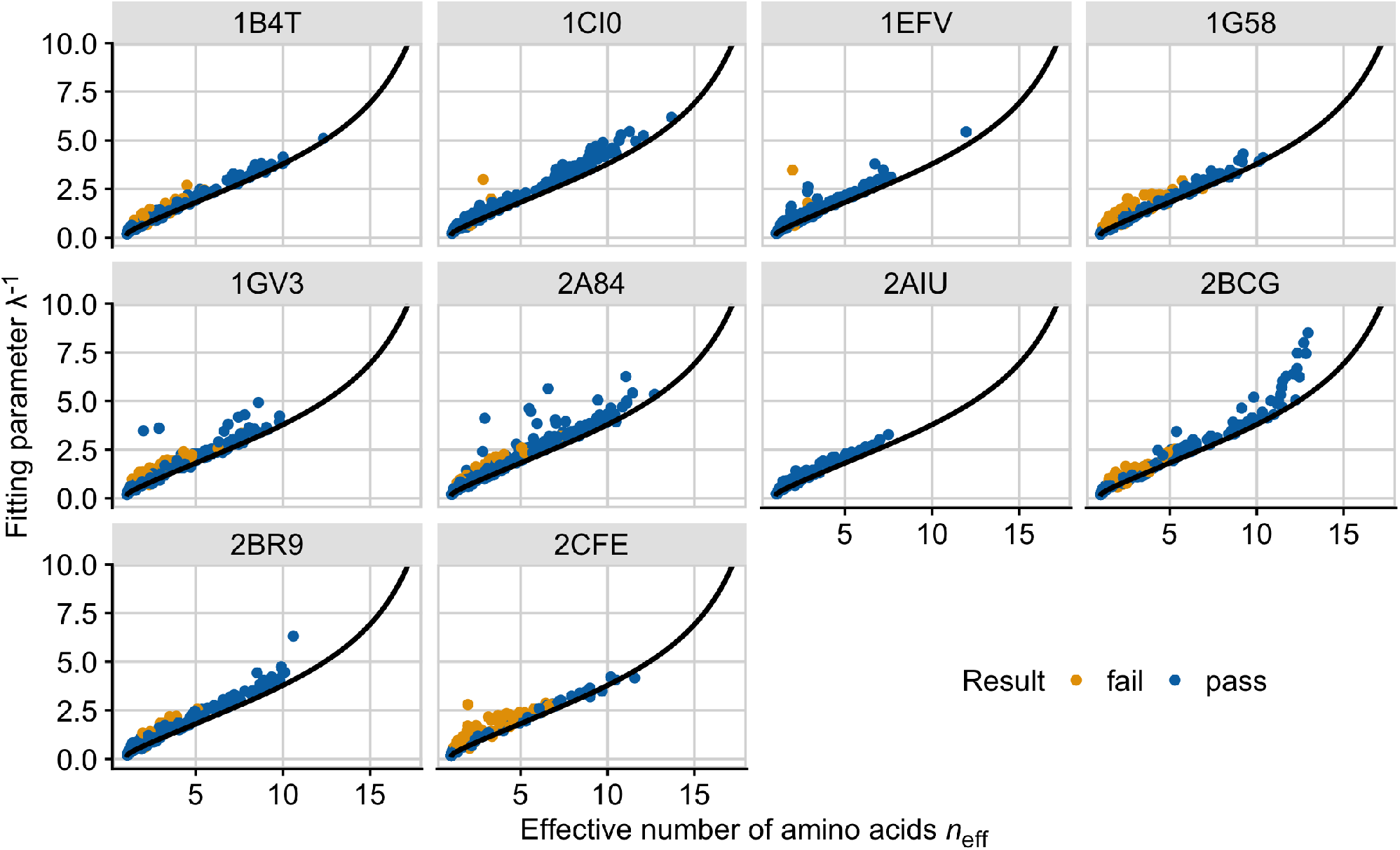
The fit of 10 empirical protein alignments to the theoretical expectation (black line, Eq. 5). Each point represents the amino acid distribution at a single site. The *x* axis represents the effective number of amino acids, *n*_eff_, calculated with Equation 1, and the *y* axis is 1*/*λ, where λ is the slope parameter from the linear regression. The title of each plot panel corresponds to the protein’s name in the Protein Data Bank. Orange points indicate a failed *χ*^2^, while blue points indicate a passed *χ*^2^ after FDR correction.

**Figure S2:**
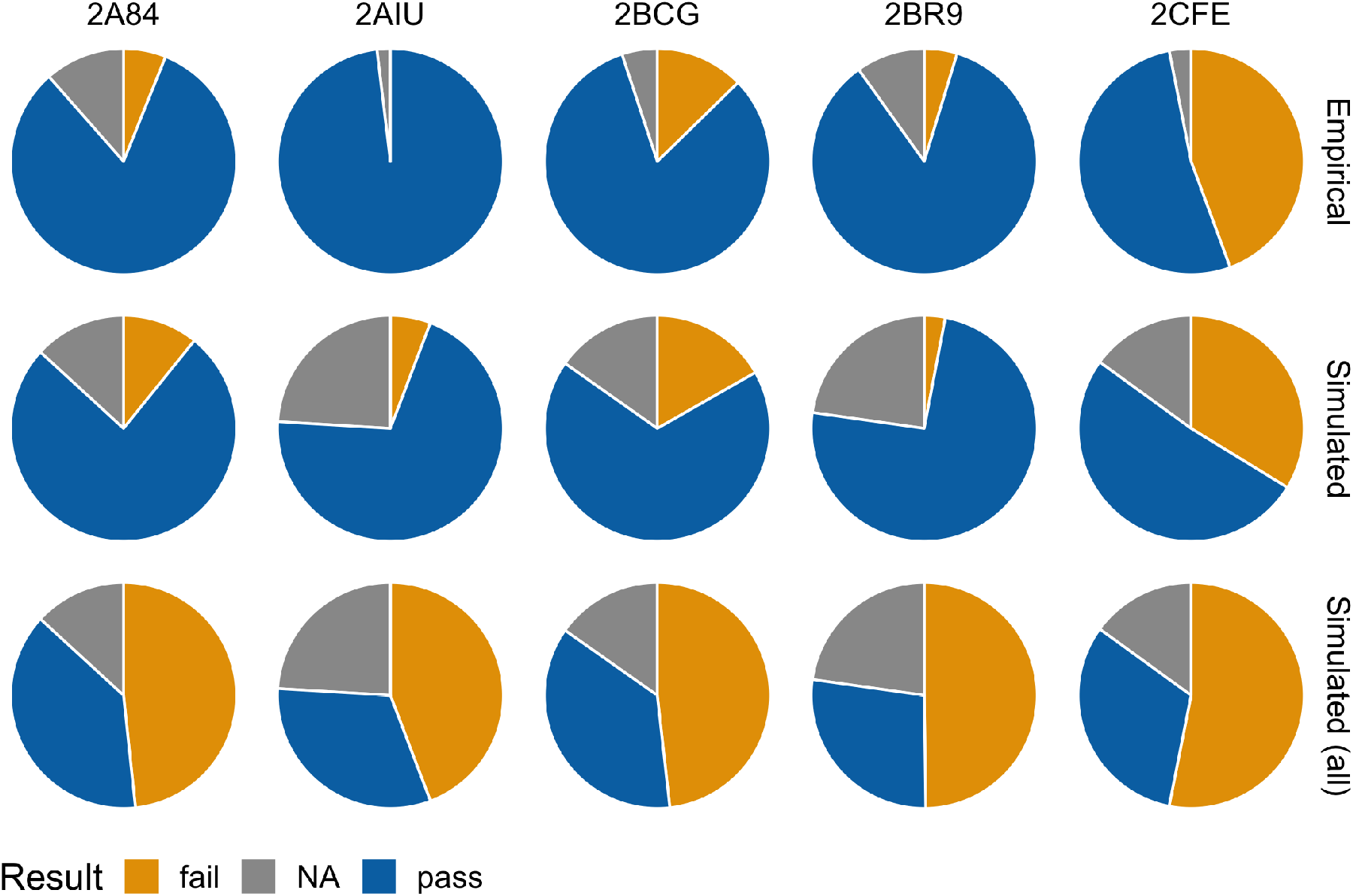
Performance of a *χ*^2^ goodness-of-fit test between the actual and estimated amino acid distributions at each site in the remaining five proteins. Analysis was performed on actual counts observed in the alignment and the counts estimated from the linear regression. Results are compared for on an empirical alignment, a simulated alignment with an equivalent number of sequences, and a simulated alignment with 500 sequences for each protein considered. A FDR correction controlled for multiple testing. Orange indicates sites that failed the *χ*^2^, while blue indicates sites that passed the *χ*^2^ and grey indicates the number of sites that could not be tested under the *χ*^2^ due to the presence of a single amino acid.

**Figure S3:**
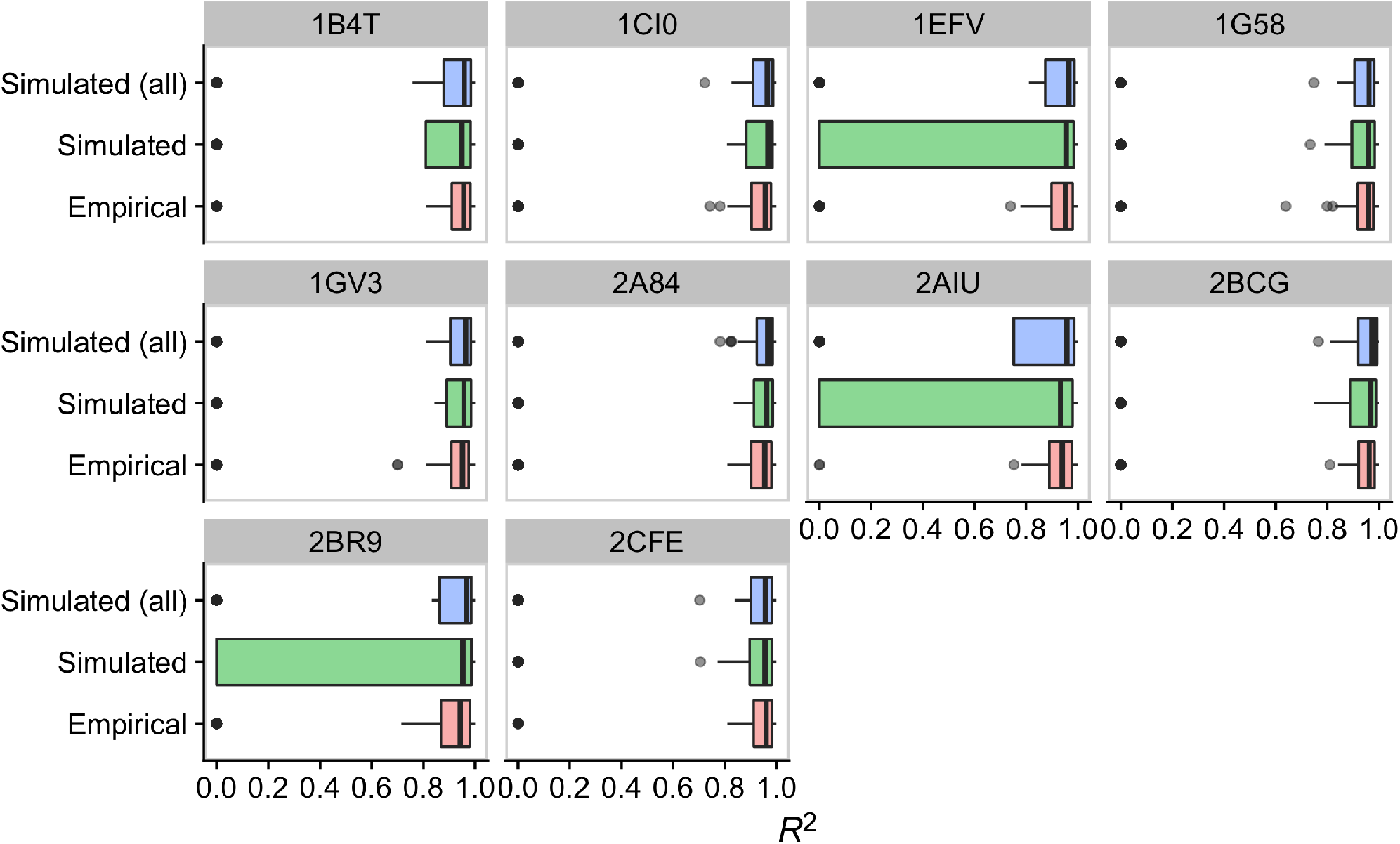
Adjusted coefficient of determination (*R*^2^) for linear regressions at sites in simulated and empirical alignments. Analysis was performed on ranked, log-transformed amino acid counts at each site. The reported *R*^2^ value comes directly from a one-parameter linear regression in R using the lm function. For each protein, the analysis was repeated on the full simulated alignment, a simulated alignment downsampled to match the size of the empirical alignment, and the full empirical alignment (see Table 1 for alignment sizes).

**Figure S4:**
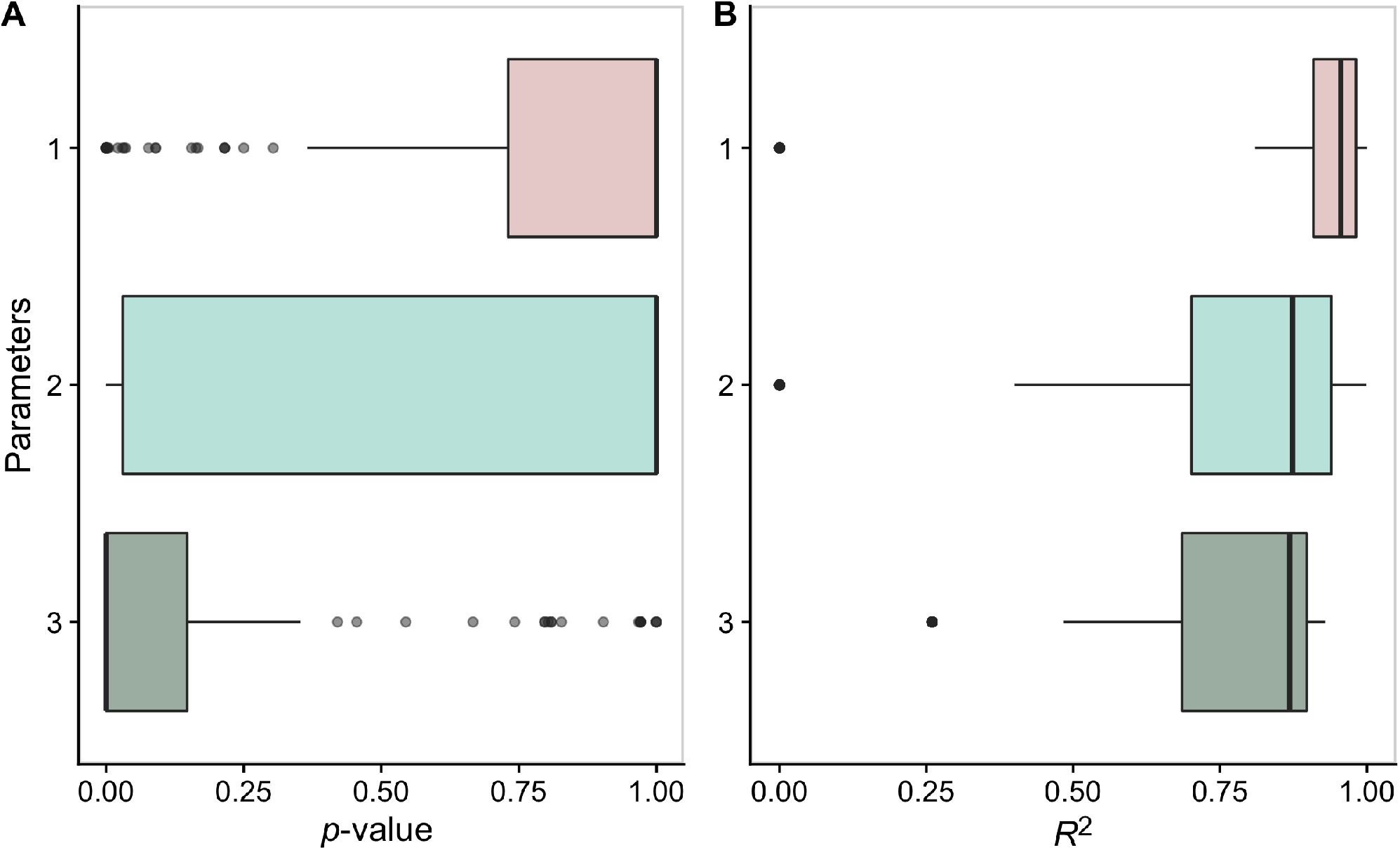
Regression performance based on the number of free parameters used to fit sites in the empirical 1B4T protein alignment. At each site, linear (one- and two-parameter) and quadratic (three-parameter) regressions were implemented and tested for their ability to reproduce the observed distribution. (A) Distribution of *p*-values from the *χ*^2^ goodness-of-fit test after FDR correction. (B) Distribution of *R*^2^ values (adjusted coefficient of determination). Each regression was fit to the ranked and log-transformed amino acids counts at a site; *χ*^2^ was tested on the actual and expected counts, while *R*^2^ comes directly from the regression fit to the transformed data.

**Figure S5:**
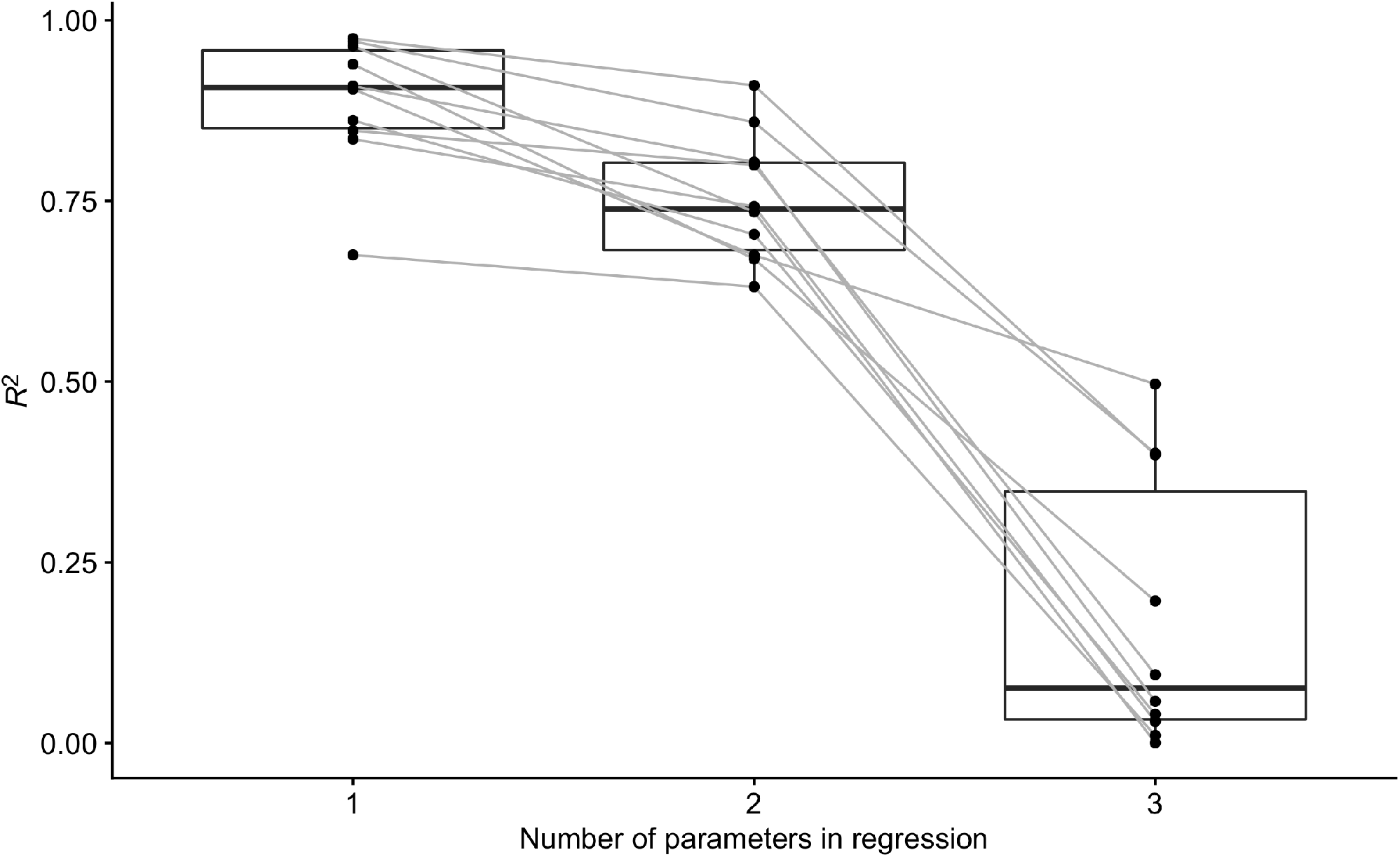
Coefficient of determination (*R*^2^) for the correlation between the effective number of amino acids (*n*_eff_) of the actual and the fit distributions, for three different regressions. Analysis shown in Figure 6 was repeated here on all 10 empirical alignments, where *n*_eff_ was calculated separately on the observed counts and the counts estimated from each regression. Each black dot represents one alignment, and lines connect alignments across regressions.

**Figure S6:**
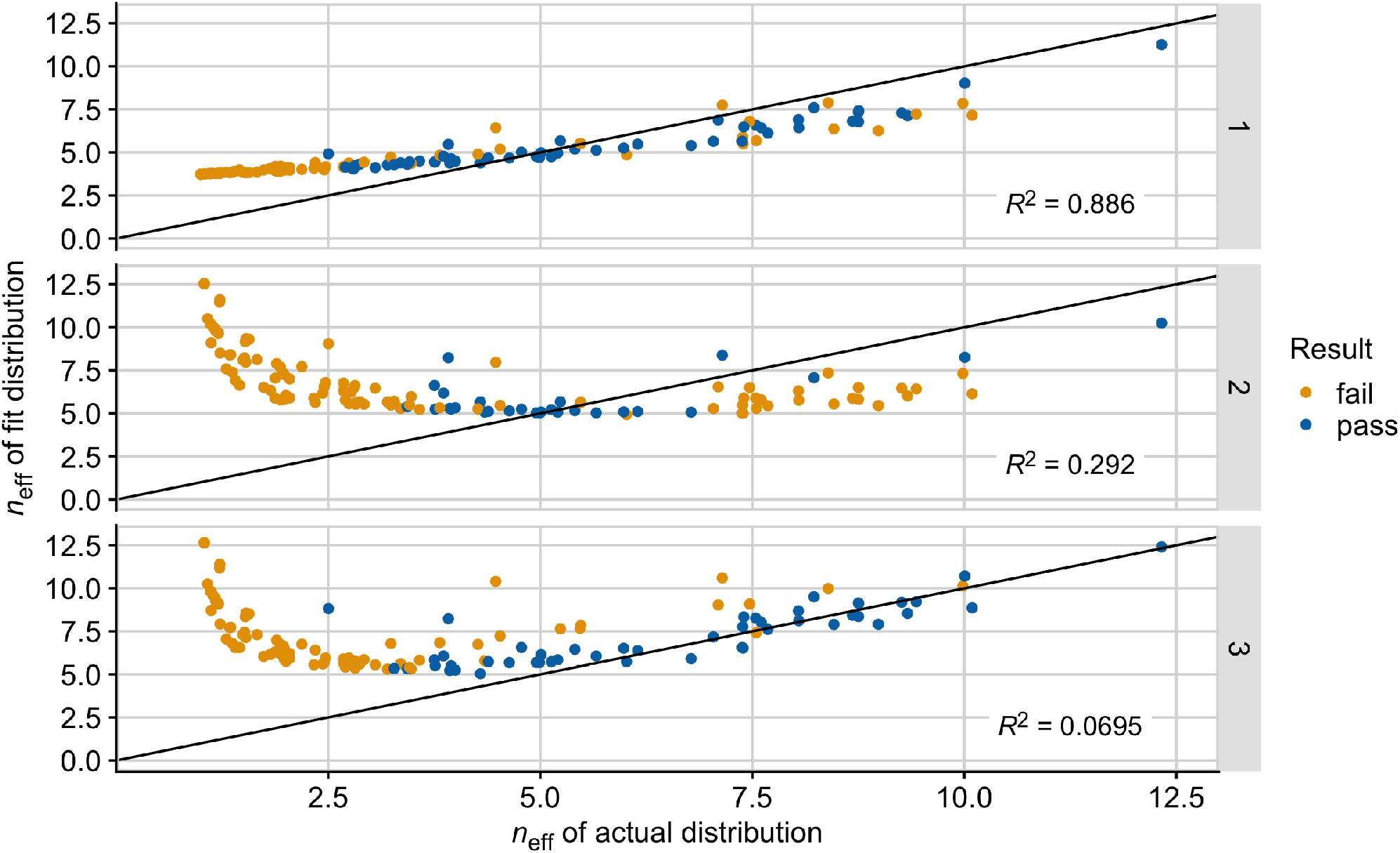
Conservation of the effective number of amino acids (*n*_eff_) in the fit distribution with modified procedures. For each site, *n*_eff_ was calculated with Equation 1 on the observed counts in the alignment and on the counts estimated from the regression analysis. Each site in the empirical 1B4T alignment was fit with either one, two, or three parameters, as indicated by the labels on the right. In Figure 6, the one- and two-parameter regressions were fit to only the amino acids present in the alignment, while a small constant was added to all counts in the three-parameter regression prior to fitting. Here, a small constant was added to all amino acids prior to fitting to ensure each regression included 20 amino acids. Additionally, the 3-parameter regression here was fit via ridge regression with tuning parameter λ_tuning_ = 0.28. The black lines represent the points for which *x* = *y*, and the *R*^2^ reported in the figure is from the correlation between *n*_eff_ in the actual and fit distribution. Colors indicate the result of the *χ*^2^ goodness-of-fit test with FDR correction.

**Figure S7:**
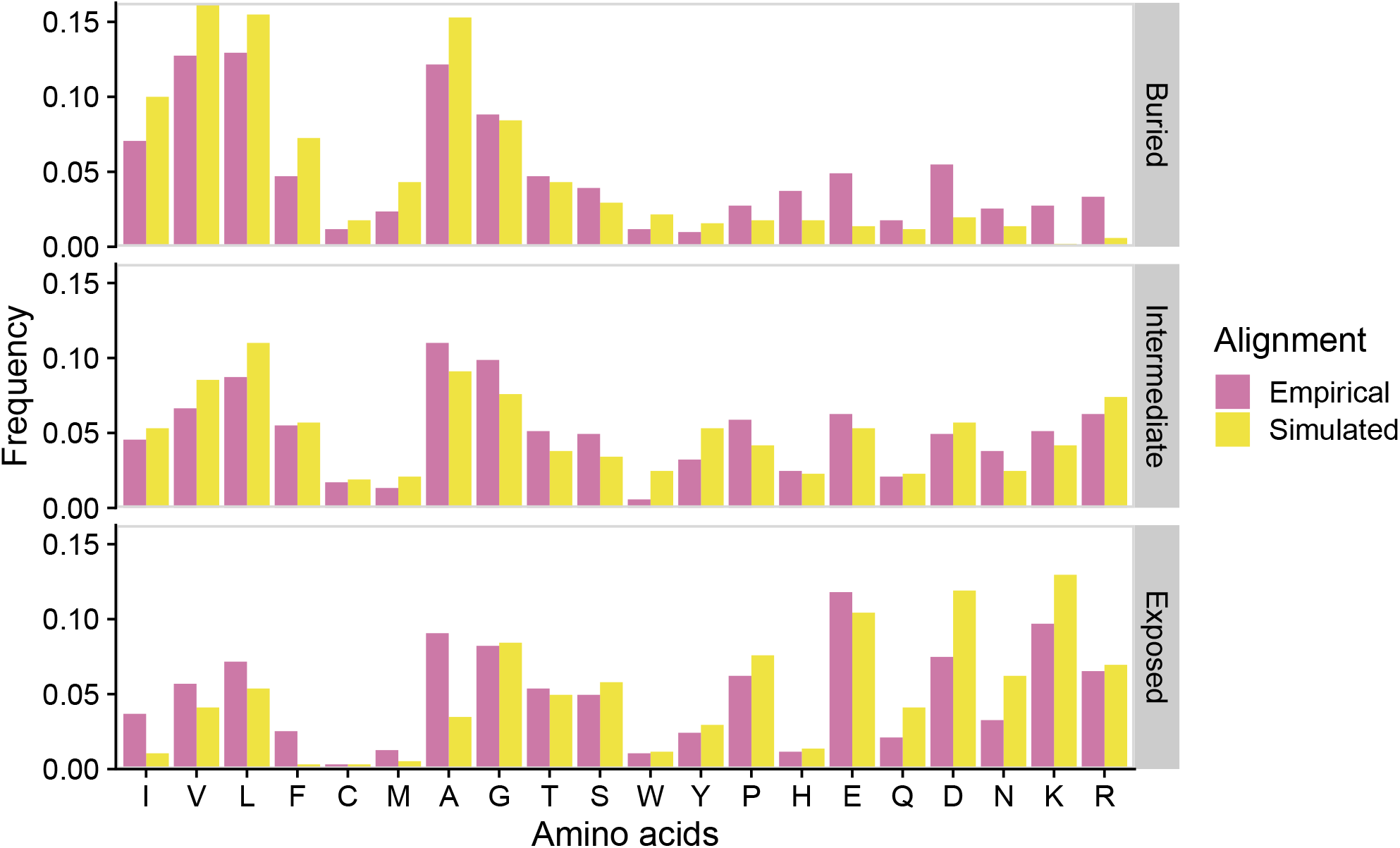
Identity of most abundant amino acid (*k* = 0) at sites based on position in the protein structure. For all 10 proteins considered, we calculated the frequency of each amino acid filling rank 0 in simulated and empirical alignments. RSA values for each site were taken from Jiang et al. (2018) and used to categorize a site’s position within the protein: buried (RSA < 5%), intermediate (5% ≤ RSA ≤ 25%), and exposed (RSA > 25%). Amino acids are listed based on hydrophobicity according to the Kyte-Doolittle scale.

**Figure S8:**
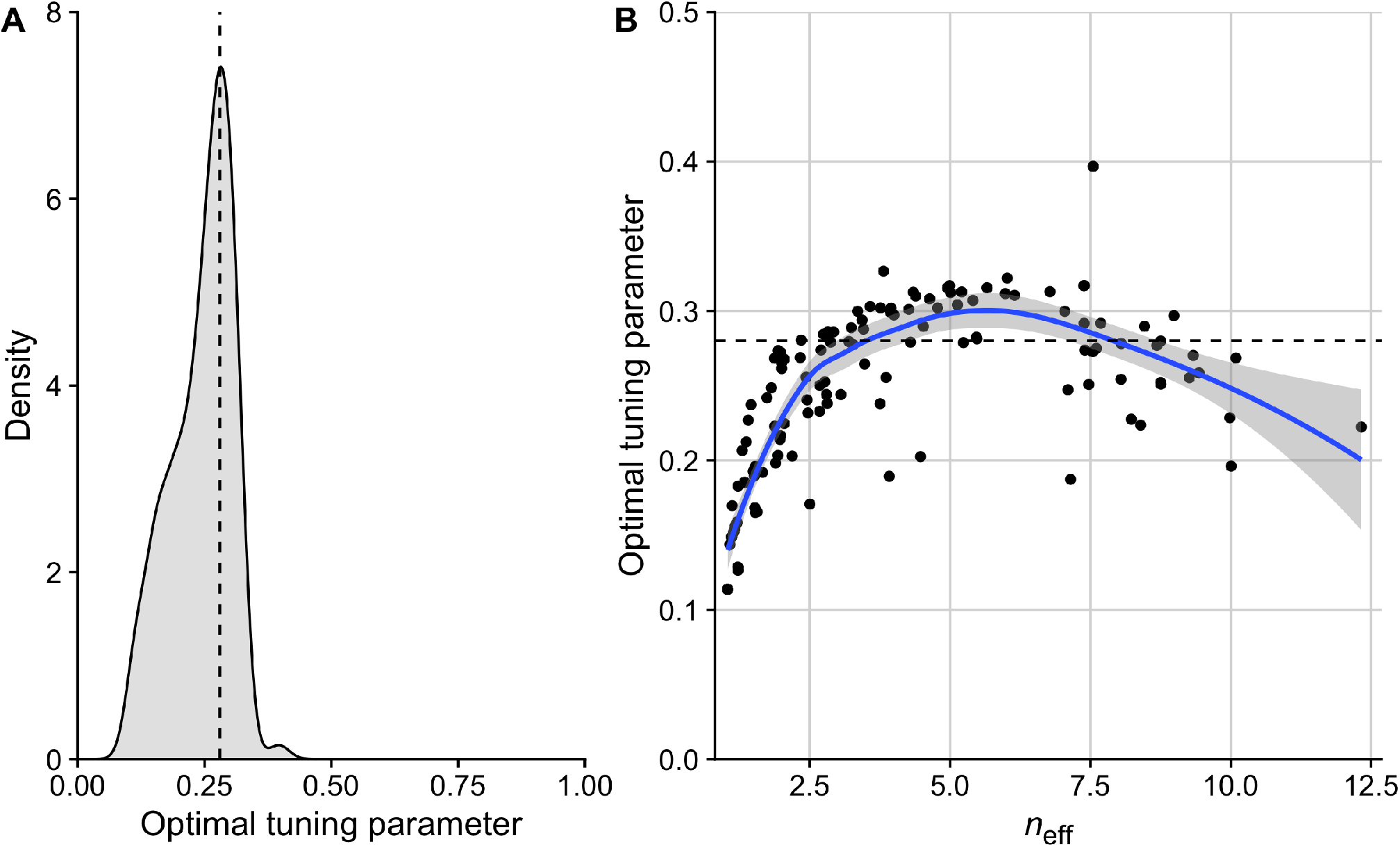
Relationship between a site’s optimal tuning parameter in a 3-parameter ridge regression and its effective number of amino acids. For sites with more than one unique amino acid observed, *k*-fold cross validation was performed in glmnet to identify the value of λ_tuning_, the tuning parameter, that gives the minimum mean cross-validation error. (A) Distribution of optimal tuning parameters. A single value was selected for use as the tuning parameter in ridge regression at all sites. (B) Effective number *n*_eff_ versus the associated optimal tuning parameter of each site. The blue line is the local polynomial regression fit with the loess function in R. In both A and B, the dashed line is the tuning parameter used at all sites (λ_tuning_ = 0.28).

